# Direct preparation of Cas9 ribonucleoprotein from *E. coli* for PCR-free seamless DNA assembly

**DOI:** 10.1101/328468

**Authors:** Wenqiang Li, Shuntang Li, Jie Qiao, Fei Wang, Yang Liu, Ruyi He, Yi Liu, Lixin Ma

## Abstract

CRISPR-Cas9 is a versatile and powerful genome engineering tool. Recently, Cas9 ribonucleoprotein (RNP) complexes have been used as promising biological tools with plenty of in vivo and in vitro applications, but there are by far no efficient methods to produce Cas9 RNP at large scale and low cost. Here, we describe a simple and effective approach for direct preparation of Cas9 RNP from *E. coli* by co-expressing Cas9 and target specific single guided RNAs. The purified RNP showed in vivo genome editing ability, as well as in vitro endonuclease activity that combines with an unexpected superior stability to enable routine uses in molecular cloning instead of restriction enzymes. We further develop a RNP-based PCR-free method termed Cas-Brick in a one-step or cyclic way for seamless assembly of multiple DNA fragments with high fidelity up to 99%. Altogether, our findings provide a general strategy to prepare Cas9 RNP and supply a convenient and cost-effective DNA assembly method as an invaluable addition to synthetic biological toolboxes.

---

Clustered regularly interspaced short palindromic repeats (CRISPR)/CRISPR associated-protein 9 (Cas9) genome-editing system is a kind of the adaptive immune systems in archaea and bacteria to defend against invasive nucleic acids from phages and plasmids^1–3^. The widely used *Streptococcus pyogenes* Cas9 (SpCas9) can be targeted to a 20 bp specific DNA sequence by an associated complementary guide RNA (gRNA), provided that a protospacer adjacent motif (PAM) of the form NRG (where R = G or A) is present^4,5^. The CRISPR-Cas9-based genetic technologies, in particular as genome-editing tools, have been extensively applied to biomedical area with great promises to revolutionize the treatment of genetic diseases^6–9^. Compared to other protein-based genome-editing technologies, including zinc-finger nucleases (ZFNs)^10^ and transcription activator-like effector nucleases (TALENs)^11^, the newly developed RNA-guided endonucleases CRISPR-Cas9 system has attracted wide attentions due to its simplicity, high efficacy, and ease of use^8,12^. CRISPR-Cas9 has been mostly used for in vivo genome excising and editing^1,3,6^, otherwise, it and its variants such as catalytically inactivated Cas9 (dCas9) have been successfully applied in high-throughput interrogation of gene functions in health and diseases^13,14^. In spite of numerous in vivo applications, recently, CRISPR-Cas9 has been employed as in vitro molecular cloning tools as well. For instance, a technique named CATCH utilized Cas9 nuclease to excise the target genome segment up to 100kilobases and ligated it to cloning vectors in a single step^15^. Besides, another technique called DASH harnessed Cas9 to effectively deplete unwanted sequence from DNA libraries prior to sequencing to reduce sequencing cost^16^. In these techniques, Cas9 functioned as a ribonucleprotein complex (RNP), in which the specific negative single guide RNA molecules (sgRNAs) bound to positively charged Cas9 enzymes. Compared to regular methods that deliver plasmids or mRNA encoding Cas9, increasing studies demonstrated that direct delivery of Cas9 RNP for genome editing in cells and animals has obvious advantages^17–20^, such as reduced off-target effects, low toxicity, high editing efficiency, etc. Therefore, a large amount of biopharmaceutical companies and research institutes have recently put greater emphasis on developing Cas9 RNP-based gene therapy medicines.

To produce Cas9 RNP, the routine strategy needs to obtain Cas9 and sgRNAs separately and then assemble them in vitro^21–23^. During the process, Cas9 is generally purified by recombinant expression from *E. coli,* whereas sgRNAs are constructed either by in vitro transcription that often requires tedious steps, or by chemical synthesis which is time consuming and expensive. Both approaches are therefore not appropriate for production of Cas9 RNPs at large scale and low cost, significantly limiting their utility for diverse applications. Here, we develop an efficient method for direct preparation of Cas9 RNP through co-expressing Cas9 and target specific sgRNAs from *E. coli.* Surprisingly, the produced RNP is super-stable with a full endonuclease activity capable of cleaving target DNA in vitro when storing at −20°C for more than half a year. The method can be used to prepare any Cas9 RNP targeting specific DNA sequences within three days, which dramatically reduces the cost of production and shows great values for large-scale and high-throughput applications.

We explored applications of the super-stable Cas9 RNP in vivo for genome editing in cells, and in vitro as molecular cloning tools, for instance, to cleave dsDNA instead of restriction endonucleases. As is known to all, the restriction endonucleases, typically have 6-or 8-bp restricted recognition sites, have been extensively used in molecular biology techniques in the past half century^24^. A plethora of powerful endonuclease-mediated assembly methods, such as Bio-Brick^25^ and Golden Gate^26^, have been developed early this century to facilitate researches in synthetic biology^27,28^. However, the restriction endonucleases have insurmountable limitations, for example, they cannot cut DNA fragments at any desired location for particular use such as seamless DNA cloning. To eliminate the needs for restriction enzymes digestion, alternative methods based on homologous recombination including sequence and ligation-independent cloning (SLIC)^29^, enzyme-independent cloning (EI)^30,31^, Gateway cloning^32^ and Gibson assembly (GA)^33^ have been developed and popularly adopted. However, these methods more or less require the use of PCR step(s) where mutations have pretty chances been introduced during either the amplification of large genes or the inverse amplification of vectors. Here, we report the development of a Cas9 RNP-based PCR-free method for seamless DNA assembly. Using the Cas9 cleavage that combines with a T5 exonuclease-assisted cloning method invented by us, we achieved simultaneous assembly of multiple DNA fragments up to 99% fidelity. The new assembly strategy termed Cas-Brick can be designed in a one-step or cyclic way, enabling efficient seamless assembly of multiple DNA fragments over 9 kilobases. Our method can facilitate researchers in the field of metabolic engineering to assemble genes from diverse organisms and construct new metabolic pathways, showing tremendous commercial potentials. Taken together, the Cas-Brick method is fully PCR-free, simple, versatile and cost-effective, enabling scarless multisegment assembly with great fidelity.

## Results

### Direct expression and purification of Cas9 RNP from *E. coli*

The functional Cas9 RNP is currently constructed by simply mixture of Cas9 and sgRNAs in vitro^23^. Usually, Cas9 enzymes were expressed and purified from *E. coli,* whereas single guide RNAs were prepared either by in vitro transcription or by chemical synthesis. However, both methods are difficult to obtain sgRNAs, which are susceptible to enzymatic degradation by ubiquitous RNase, in sufficient amounts and high quality. Previously, researchers successfully achieved production of mature ribonucleoproteins in heterogeneous expression systems^34,35^. For instance^34^, they introduced two plasmids to co-express Cas9-crRNA (representing short CRISPR RNA) complex, in which one plasmid encoded Cas9 enzyme and the other encoded Cas1, Cas2, Csn2, SP1 and transencoded small RNAs (tracRNAs) in addition to short CRISPR RNAs (crRNAs). This expression system has many defects, such as complexity, low yield of Cas9-crRNA (1.5 μg from 1 L *E.coli* culture) complex and unsatisfied Cas9 nuclease cleavage activity, significantly limiting its widespread uses. Herein, we attempt to establish a new co-expression system to efficiently prepare functional Cas9 RNPs. To achieve the goal, we generated a plasmid based on an engineered cold-shock vector pCold I^36^ (Fig. 1, Fig. S1) which prevents leaking expression of Cas9 at 37°C which is for *E. coli* grown, while guarantees sustainable expression of Cas9 at 16°C after adding IPTG. Meanwhile, sgRNA molecules were abundantly transcribed in vivo by *E. coli’s* own RNA polymerases using a T7 promoter. According to the design, we hypothesized that the newly synthesized Cas9 and transcribed sgRNAs would be spontaneously self-assembled in *E. coli* cells to form complete Cas9/sgRNA complexes.

**Fig. 1.**
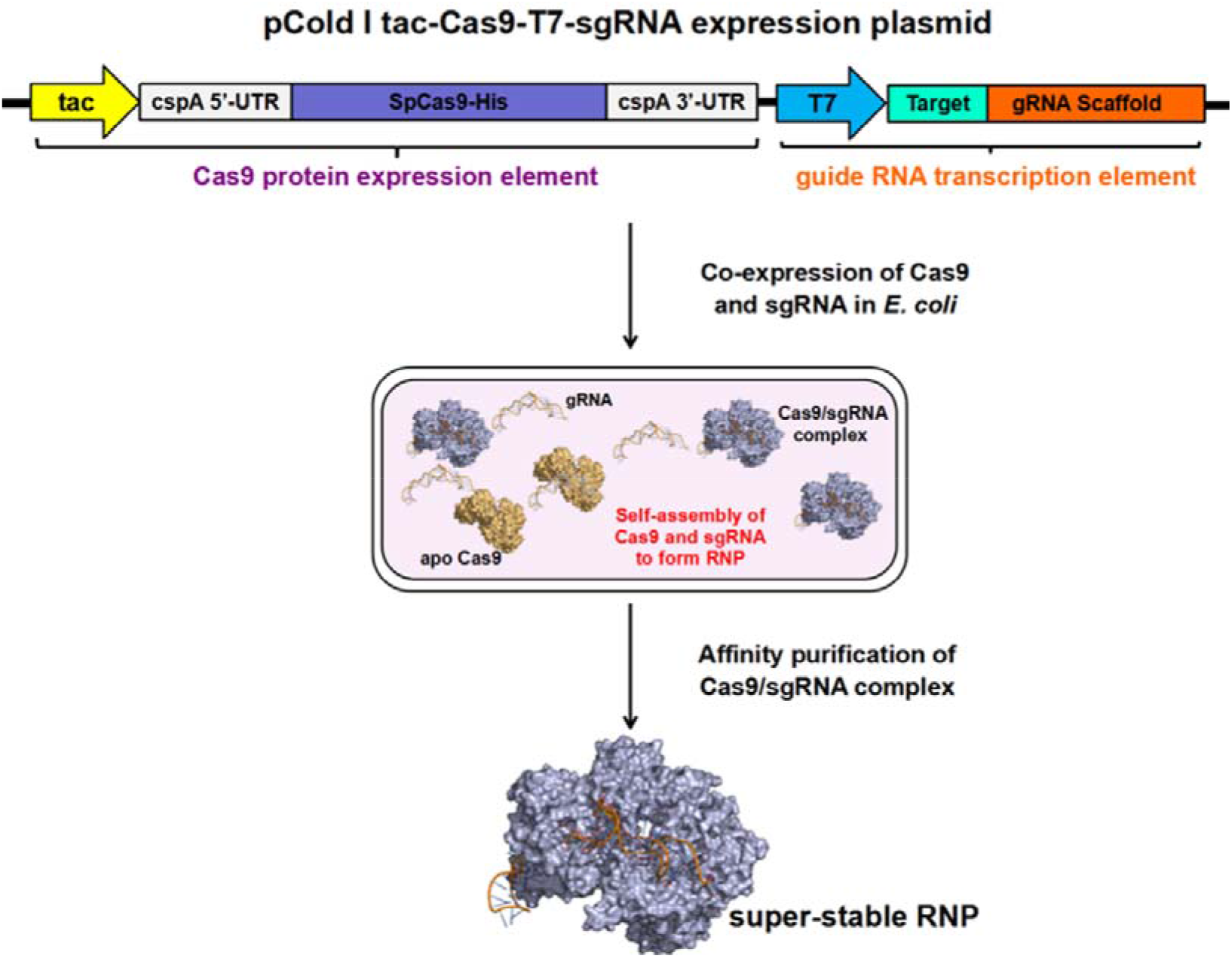
An engineered cold-shock pCold I plasmid was harnessed to achieve co-expression of *SpCas9* and sgRNAs in *E. coli.* The newly synthesized apo-Cas9 (yellow) and sgRNA (helical) are supposed to self-assembly to form a super-stable Cas9/sgRNA complex (blue) that can be readily purified.

To verify the hypothesis above, we firstly purified such Cas9 RNP by affinity purification using Ni-NTA column. The RNP showed an endonuclease activity capable of recognizing and cleaving target dsDNA in vitro (Fig. 2a). There is no needs to add RNase inhibitors during the whole purification process. Conversely, a sufficient amount of RNase inhibitors are required to prevent degradation of sgRNAs when using traditional methods to produce Cas9 RNP. To the best of our knowledge, it is by far the simplest and cheapest way to produce Cas9 RNP. More surprisingly, the obtained RNP is super stable that maintained full enzyme activity when storing at 4°C for three months, or at −20°C for up to half a year in absence of RNase inhibitors (Fig. 2a), leading to significant commercial development in future. To explain the high stability of RNP, we proposed that the sgRNAs transcribed in vivo might bind to pre-formed Cas9 tightly, thereby were protected from nuclease-mediated degradation. Another explanation is that the low temperature for Cas9 expression may reduce RNase activity in *E. coli.*

**Fig. 2.**
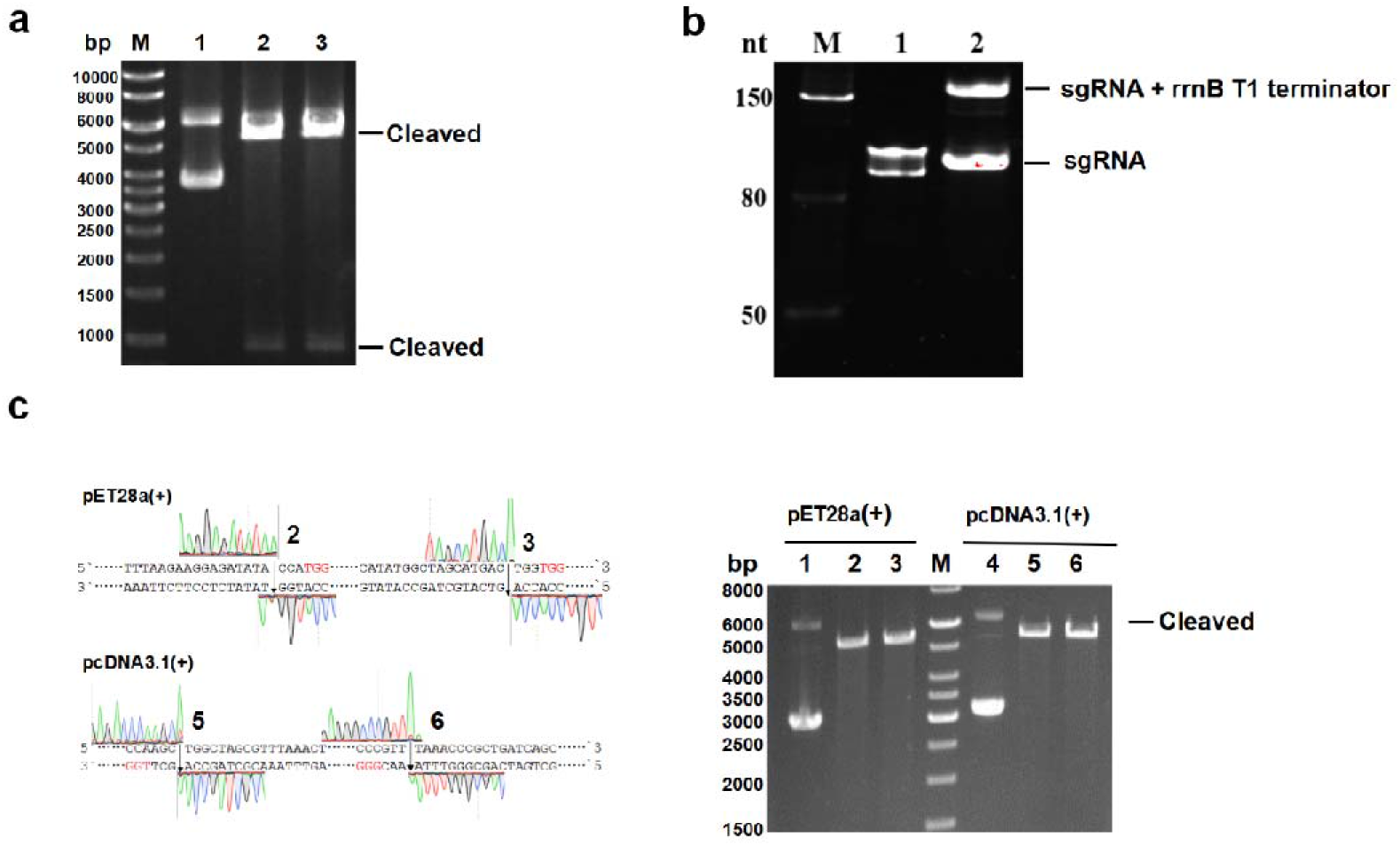
Characterization of Cas9 RNP.**a.** Activity of Cas9 RNP on cleaving the plasmids (lane 1) after storage at 4°C for 90 days (lane 2) and at −20°C for 180 days (lane 3). We introduced two cleavage sites with the same recognition sequence, producing a DNA fragment at 1000 bp after cleavage by Cas9 RNP. **b.** The RNA electrophoresis gel (8% polyacylamide) of sgRNA produced by in vitro (lane 1) transcription, or by in vivo (lane 2) transcription. **c.** Cleavage of plasmids including pET28a(+) (lane 1) and pcDNA3.1(+) (lane 4) at MCS using two Cas9 RNPs. The cleavage sites were indicated with arrows and numbered (left panel) corresponding to the lanes on the right.

To characterize Cas9 RNP made in this way, we treated it with proteinase K to digest Cas9 enzymes followed by extraction of incorporated RNA molecules. The RNA electrophoresis gel assays resulted in two homogeneous bands, namely one band at 100 bp that contains RNA sequences identical to the corresponding sgRNA transcribed in vitro (Fig. 2b), as well as the other band at 170 bp which includes sgRNA plus additional rrnB T1 terminator sequences (Fig. S2). It has been proven that 3’-extra RNA sequences of gRNA unaffected nuclease activity of Cas9^37^. In this study, we found that the Cas9 enzyme plus 170 bp sgRNAs extracted from Cas9 RNP in vitro, with 3’-extra rrnB T1 terminator sequence, has a full nuclease activity (Fig. S3). Moreover, RNA-seq experiments verified that the sequence of incorporated RNA is in agreement with design (Fig. S4), proving that sgRNA can be precisely in vivo transcribed.

### Specific cleavage of vectors by Cas9 RNPs instead of restriction endonucleases

Restriction enzymes are essential genetic tools for recombinant DNA technology that have revolutionized modern biological research since the early 1970s^38^. So far, there are over 250 commercially available restriction endonucleases for routine uses in thousands of laboratories around the world^39^. Most of them only recognize short DNA sequences (typically ~6-or 8-bp), which limits their applications in particular use such as seamless DNA cloning. Recently, Wang et al have successfully employed CRISPR-Cas9 enzymes as artificial restriction enzymes (AREs) combined with Gibson assembly to accomplish seamless DNA cloning^40^. Yet, it is difficult to prepare AREs in sufficient amount with low cost, significantly challenging their wide applications instead of restriction enzymes. Herein, our aforementioned method addresses the challenge, enabling large-scale production of super-stable Cas9 RNP.

Typically, Cas9 recognizes a ~ 20 bp target sequence with a required downstream NRG (where R= G or A) PAM and induces a site-specific double strand break (DSB)^5,41,42^. Thus, we firstly searched all PAM sites at multiple clone sites (MCS) in the common used vectors, such as pET28a(+) and pcDNA3.1(+). For example, the vector pcDNA3.1(+) has twelve restriction enzyme sites at MCS, while it has nineteen PAM sites in the same region. Even better, we are able to simply introduce new PAM sites at any desired location in the vectors. It is therefore more than enough to generate sufficient Cas9 RNPs instead of restriction enzymes to cut the vectors. The multiple PAM sites can also facilitate researchers to choose specific sites for molecular cloning in certain situations that restriction enzymes were forbidden. For instance, we cannot use a restriction enzyme if its recognizing sequence is present both in cloning DNA fragment and vector. Nevertheless, we do not have to worry about this issue when using Cas9 RNP, since there is a pretty low probability that the same 20 bp recognizing sequence both exists in cloning DNA and vector.

To determine in vitro nuclease activity of Cas9 RNP, we cleaved plasmids (Fig. 2c) by different concentrations of RNP to generate blunt ends. Typically, 300 ng plasmids were cleaved in less than 30 minutes at 37°C when 200 ng of RNPs were added. It has to be pointed out that the concentration of RNP is determined by Cas9 protein instead. By the method, over 20 mg Cas9 RNPs (sufficient for > 10^5^ reactions) were generally produced from 1 L of *E. coli* LB culture, which is over 10^4^ fold than the yield of Cas9 RNPs in the previous work^34^. At present, we have established an efficient platform for readily producing highly active Cas9 RNPs as AREs for common molecular biology uses.

### Single Cas9 RNP-based seamless DNA assembly

Nowadays, Gibson assembly (GA) (Fig. S5) has become one of the most broadly used methods for in vitro DNA assembly^33^. Though GA is fast and versatile, it is very expensive because it requires three enzymes including T5 exonuclease, Phusion DNA polymerase and Taq DNA ligase. Meanwhile, enzyme-independent cloning (EI) methods^43,44^ have been developed capable of assembling DNA fragments without any enzyme at low cost. In a typical EI cloning (Fig. S5), the insert and linear plasmid assembled at over 15 bp nucleotides long homology regions through an *E. coli* repair or recombination event that is not fully understood. In this work, we employed EI cloning or an simplified GA termed T5 exonuclease assisted cloning invented by us (Fig. S5) to fulfill seamless assembly of DNA fragment(s) derived from Cas9 RNP cleavage. In accordance with previous reports^43–45^, we found that the efficiency of EI method drops sharply when the number of gene fragments is over three. So, the T5-assisted cloning is preferable when assembling three or more gene fragments. This method is pretty easy to follow since it only needs to roughly mix T5 exonuclease, gene fragment(s) and linear vector on ice for five minutes followed by transformation. The gaps between the gene fragment(s) and linear vector will be repaired and eventually ligated to form a complete plasmid by enzymes in *E. coli.* Thus, the T5-assisted cloning is more cost-effective and convenient than GA, which has become a huge hit with the laboratories in our university.

The work flow of a typical three-fragment assembly was shown in Fig. 3. Accordingly, only a single Cas9 RNP was adopted to cleave donor and acceptor plasmids. To simply select monoclonal colonies from phenotypes, we choose three gene makers encoding green fluorescence protein (GFP), red fluorescence protein (RFP) and kanamycin-resistant protein (KR). If they were all correctly assembled, the three proteins will be formed resulting in orange colored colonies under ultraviolet light on culture plates (Fig. 3) with both kanamycin and ampicillin. The assurance of clones can be further verified by direct DNA sequencing. It should be noted that the 6 bp protruding end of gene fragments, a product of RNP cleavage, will be depleted during the recombination in *E. coli.* Thereby, we successfully achieved scarlessly assembly of target DNA fragments. The results of EI and T5-assisted cloning with lengths of 20 bp, 30 bp and 40 bp homologous overlaps were summarized in Tab. 1, revealing that the number of colonies rose remarkably along with increasing lengths of overlaps. More importantly, almost 100% fidelity can be achieved when using T5-assisted cloning with over 30 bp homologous overlaps. Collectively, considering both recombinant efficiency and fidelity, we eventually choose 40 bp homologous overlaps in the following assembly experiments unless otherwise noted. Statistically, 150 c.f.u./ng cloning efficiency was observed for a transformation experiment.

**Fig. 3.**
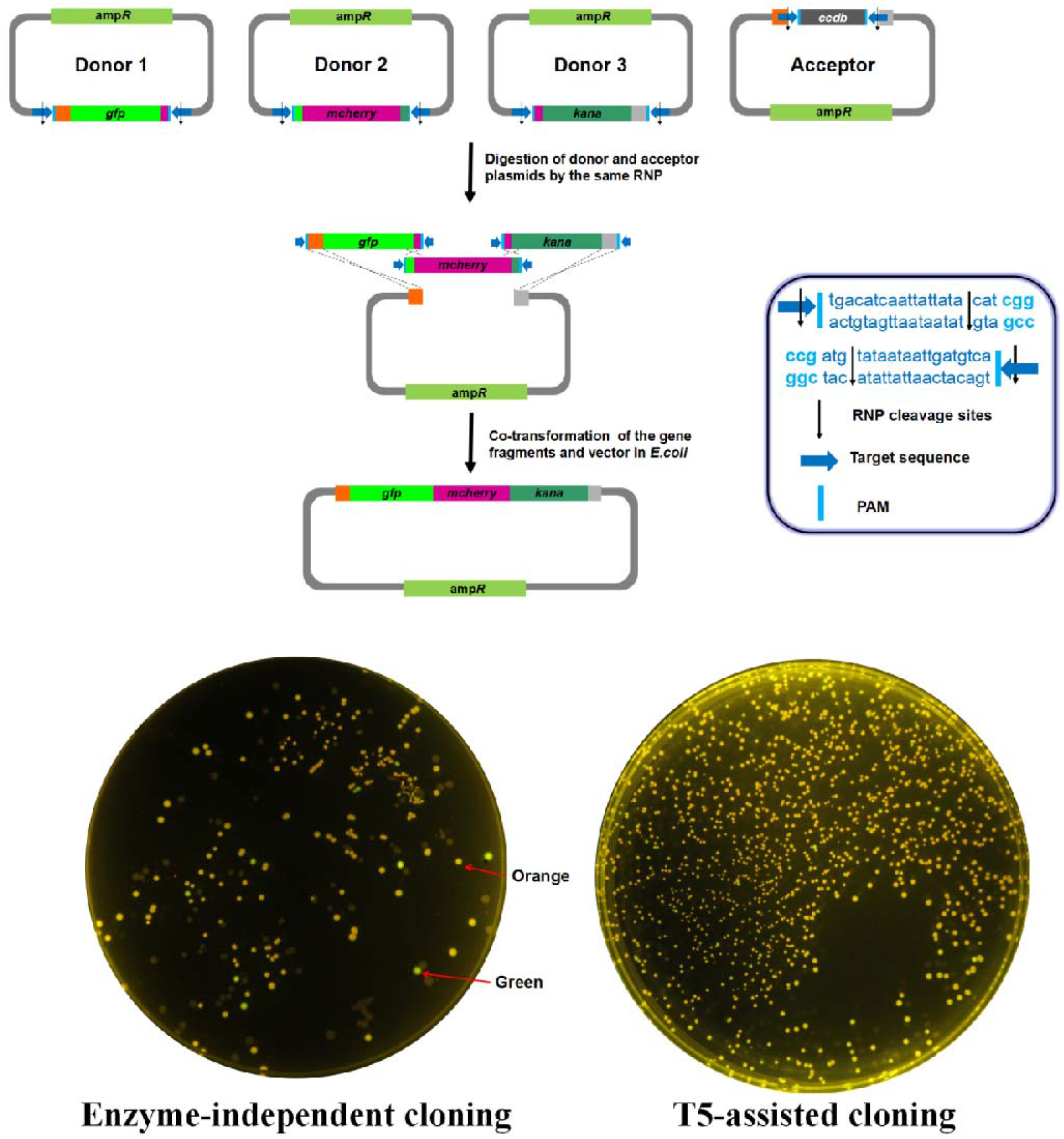
The top panel shows the flow diagram of a representative three-fragment assembly. Accordingly, three donor plasmids and one acceptor plasmid were cleaved by the same Cas9 RNP, generating three DNA fragments encoding GFP (green), Mcherry (red) and Kana (dark green) as well as a linear accept vector. The insert shows PAM and RNP cleavage sites. The bottom panels show the assembly results by harnessing an enzyme-independent cloning or T5-assisted cloning method with a 40 bp homologous overlaps involved.

**Tab. 1:** A three-fragment DNA assembly with different homologous overlaps.

| Homologous overlaps (bp) | Enzyme-independent cloning |  | T5-assisted cloning |  |
| --- | --- | --- | --- | --- |
|  | Clones<sup>a</sup> | Fidelity<sup>b</sup> | Clones | Fidelity |
| 20 | 148/365 | 40.5 ± 2.1% | 472/814 | 57.6 ± 2.5% |
| 30 | 137/324 | 42.3 ± 4.2% | 1868/1871 | 99.8 ± 0.2 % |
| 40 | 538/731 | 73.6 ± 11.2% | 1993/1994 | 99.7 ± 0.2 % |

a. The data was given as “positive clones/total clones” by counting the number of clones from three independent experiments for each methods. The positive clones appeared orange color on the culture plates having both kanamycin and ampicillin. b. The fidelity was calculated from observed phenotypes. The accuracy of clones were further verified by DNA sequencing through sending twenty clones of each plate.

### Cas-Brick cyclic assembly

Bio-Brick standard was proposed early this century which employs *Eco*R I and *Xba* I cutting sites as the prefix sequence and *Spe* I and *Pst* I as the suffix sequence for hierarchical assembly^25^. However, it cannot be used for the construction of in-fusion proteins, due to an 8-bp scar between parts exist. Several other assembly methods therefore have been developed to solve the problem, such as BglBrick^46^, iBrick^47^ and the newly developed C-Brick^49^. Here, we develop an efficient method termed Cas-Brick to fulfill cyclic assembly of multiple gene fragments with high fidelity.

Taking a nine-fragment assembly for example (Fig. 4), four pairwise Cas9 RNPs were adopted (Fig. 4a) to cleave two acceptor plasmids with distinct resistance, and to cut donor plasmids in three ways at demand. Initially, we carried out three paralleled individual three-fragment assembly in the first round. Next, we cleaved three newly synthesized DNA parts using pairwise RNPs and subsequently cloned them to another pre-formed linear vector in the second round, accomplishing seamless insertion of a 9 kilobases large DNA fragment at fidelity over 95%. As shown in Fig. 4c, the correct clones showed red color under white light on behalf of production of astaxanthin. Meanwhile, they also appeared green color under ultraviolet light on behalf GFP. Collectively, this method can significantly facilitates researchers in the field of metabolic engineering to screen wanted colonies. The accuracy of selected colonies was further verified by DNA sequencing by sending the ones with correct phenotypes. Additionally, the routine of Cas-Brick assembly can be designed in multiple-round cyclic, making it suitable for multisegment assembly of large DNA constructs.

**Fig. 4.**
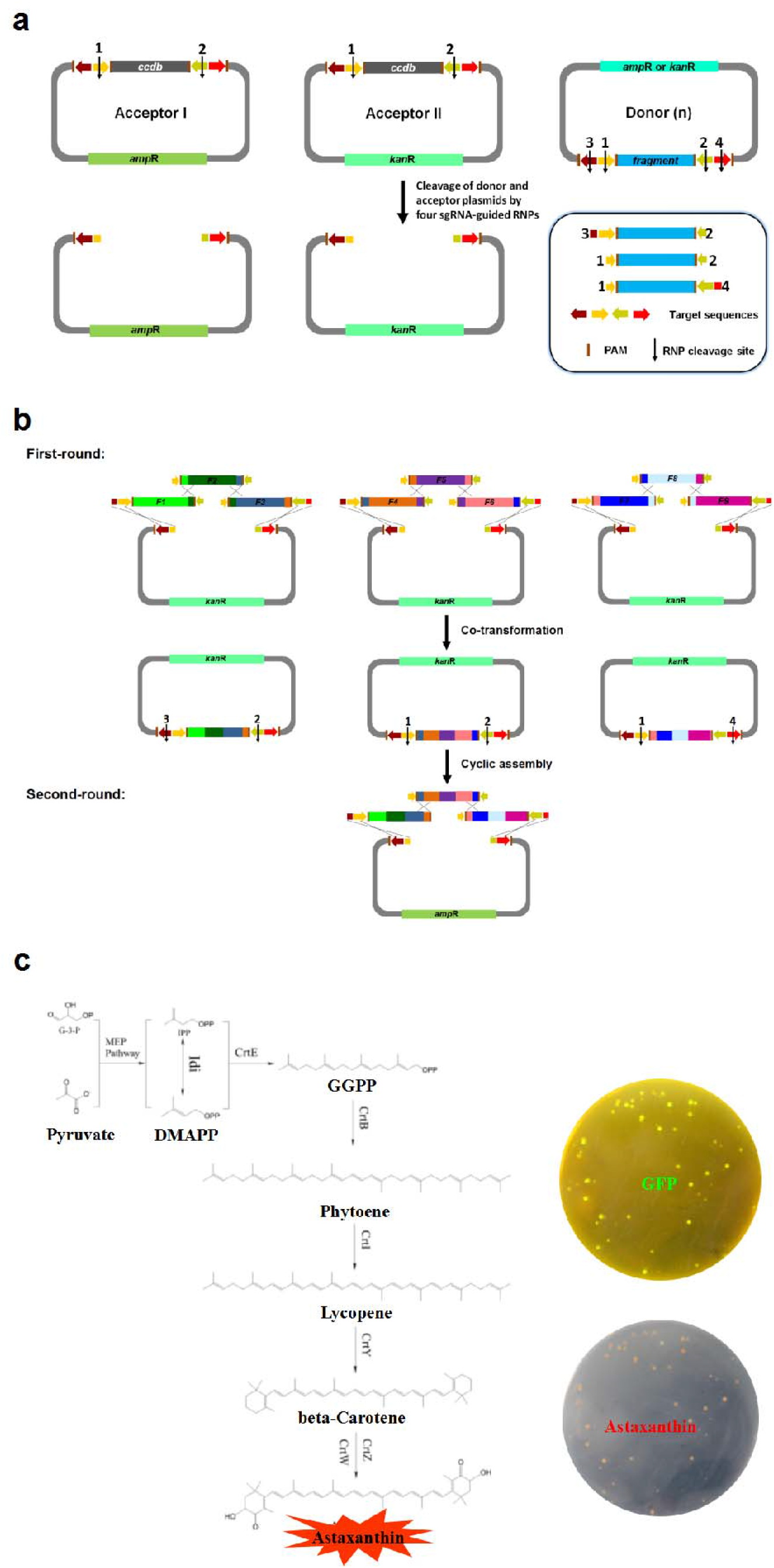
Cas-Brick method for cyclic mutisegment assembly. (a) In a standard Cas-Brick, four pairwise Cas9 RNPs were employed to cleave acceptor I (ampicillin resistance), acceptor II (kanamycin resistance), as well as donor to generate linear vectors and DNA fragments. The insert panel shows three distinct cleavage ways. (b) Assembling nine individual gene fragments by two rounds of Cas-Brick. The linear vectors with kanamycin and ampicillin resistance were alternately used in each cycle. F1-F9 represent nine DNA fragments coding GFP, CrtE (geranylgeranyl pyrophosphate synthase), CrtI (phytoene dehydrogenase), CrtB(phytoene synthase), Idi (isopentenyl-diphosphate delta-isomerase), CrtY (lycopene cyclase), CrtZ (β-carotene hydroxylase), CrtW (β-carotene ketolase), LacI, respectively^48^. Among them, the correct expression of F2 to F8 will produce a series of enzymes leading to the synthesis of astaxanthin in *E. coli.* (c) The biosynthetic pathway for synthesis of red colored astaxanthin. The correct colonies can be easily screened by phenotype, because they will appear red under white light and green under ultraviolet light.

## Discussions

The programmable nuclease Cas9 has been shown to mediate editing of disease-associated alleles in the human genome, facilitating new treatments for many genetic diseases^6,9^. Traditionally, Cas9 can be produced in cells by delivering the corresponding coding sequences as either DNA or mRNA^42^. Recently, direct delivery of pre-formed Cas9 RNP instead has been proven as a promising method for genome editing with advantages such as high editing efficiency, reduced off-target effects, rapid clearance in vivo, etc^17–20^. The current strategy to construct RNP is by mixing Cas9 and one or more sgRNAs in an appropriate buffer. However, it is generally expensive to prepare Cas9 RNPs in sufficient amounts by this method. Here we describe an efficient method to directly prepare RNP from *E. coli* cells. Initially, we harnessed an engineered cold-shock expression plasmid to co-express Cas9 enzymes and sgRNAs in *E. coli.* According to the design, the newly synthesized Cas9 and sgRNAs would spontaneously self-assemble in vivo to form complete Cas9 RNPs. To characterize them, we performed RNA electrophoresis gel assays which indicate that sgRNAs transcribed in vivo were homogeneous, containing the full designed RNA sequences which is further verified by RNA-Seq. We hypothesize that the simultaneous incorporation of sgRNAs into newly synthesized Cas9 in vivo might help Cas9 folding into a more stable conformation. Remarkably, we found that the produced Cas9 RNP has a superior stability that maintained full enzyme activity without RNase inhibitors when stored at −20°C for up to half a year.

Nowadays, it has been proven that injection of Cas9 RNP into tissues and organs is a promising strategy for the treatment of genetic diseases^50^, however, one of the most challenges remaining is that RNP must be stable enough to survive degradation in organisms^51^. Limited protein lifetime will require delivery of higher doses of Cas9 into the patient, resulting in poor target editing. Conversely, delivering a RNP with improved thermostability showed an increased lifetime in human plasma^51^. Our data demonstrated that delivering the super-stable Cas9 RNP made in this work has a profound genome-editing efficiency in mammalian 293T cells (Fig. S6). More experiments are ongoing towards other cell lines, tissues and animal models to unravel the detailed molecular basis. Recently, CRISPR-based screening of genomic DNA has enabled studying both genes and non-coding gene regulatory elements, but construction of high-throughput screening platforms using Cas9 RNP libraries have not yet been established, largely due to production of Cas9 RNP is time-consuming and expensive by current methods. To address the challenge, we provide here an efficient method to prepare Cas9 RNP at large scale and low cost. Strikingly, the method can be applicable to produce other programmable nuclease RNP such as Cpf1 (Cas12a)^52^ and C2C2 (Cas13a)^53,54^, thereby showing extensive commercial potentials in future. We have applied patents relating to the method, and believe that popularization of them will enable establishing high-throughput gene screening platforms as well as developing RNP-based reagents and vaccines.

In the second part, we firstly constructed a series of Cas9 RNPs to effectively cleave plasmids instead of broadly used restriction endonucleases. By far, we have completely replaced common restriction enzymes with the corresponding Cas9 RNPs for regular molecular cloning in our lab. In a word, researchers can utilize our method to produce any AREs according to their own requirements, which is especially useful for certain cases that restriction enzymes were not allowed. The easy-to-use Cas9 RNP is capable of cleaving any target dsDNA, unrealizing potential to revolutionize the practice of molecular biology well beyond genome editing. There are recently increasing needs for the precise DNA assembly with fast development of synthetic biology. The holy grail of DNA assembly was proposed to be a method that allows scarless, sequence independent, multi-fragment assembly of large DNA fragments at high efficiency and fidelity^45^. But to date, none of assembly techniques has been able to fulfill the ideal and each method has its own merits and defects that were summarized in Tab. 2. Here, we report a PCR-free DNA assembly method called Cas-Brick with enormous progress close to the ultimate goal. We harnessed Cas9 RNP to cleave plasmids for generation of DNA fragments and linear accepted vectors. The assembly routine can be easily designed at demand in a one-step or cyclic way. Combined with a T5 nuclease-assisted cloning method, we achieved seamless assembling multiple DNA fragments up to 9 kilobases at high fidelity close to 100%. The Cas-Brick is principally multiple-round cyclic, and the assembly efficacy is dependent on transformation efficiency of large DNA fragments. Our method also provides a simple platform for screening colonies, largely facilitating researchers in the field of metabolic engineering. Strikingly, T5 nuclease-assisted cloning method is more cost-effective and convenient compared to the broadly used Gibson assembly^30^. All in all, our Cas-Brick will be an invaluable addition to the synthetic biological toolboxes.

**Tab. 2.** Comparison of multisegment DNA assembly methods

|  | Gateway | Bio-Brick | Golden-gate | SLIC | Gibson | Yeast HR | LCR | Cas-Brick |
| --- | --- | --- | --- | --- | --- | --- | --- | --- |
| Days to Completion<sup>a</sup> | 4 | 3 | 3 | 3 | 3 | 5 | 3 | 3 |
| PCR Dependent | no | no | no | yes | yes | yes | yes | no |
| Scarless | no | no | yes<sup>b</sup> | no | yes | yes | yes | yes |
| Restriction Site Dependent<sup>c</sup> | no | yes | yes | no | no | no | no | no |
| Construction Over 30 kb<sup>d</sup> | no | no | no | no | yes | yes | yes | yes |
| Costs<sup>e</sup> | \$\$\$\$ | \$\$ | \$\$ | \$\$ | \$\$\$ | \$\$ | \$\$ | \$ |

a. Time in days for a three-segment assembly (two inserts plus a vector) starting with PCR segments or restriction fragments and ending with an assembled plasmid purified from *E. coli*.
b. Golden-gate can be quasi-scarless, using type IIs restriction enzymes which cut at sites outside their recognition sequences, such as *Bsa* I (GGTCTCN^NNNN) and *Bpi* I (GAAGACNN^NNNN).
c. Limited by restriction enzyme recognition sites. Sites found within segments need to be mutated before assembly.
d. Constructions over 30 kb will result in adequate efficiency and accuracy.
e. The costs here include both up front costs such as purchasing vectors and the time course to construct plasmids, and the consumable supply costs associated with DNA assembly that would change depending on scale.

## Methods

### Construction of plasmids

The expression plasmid was constructed based on pCold □(Takara) plasmid by replacing the original cspA promoter with a tac promoter, and by replacing the f1 ori with a T7 promoter. The spCas9 gene from pET-Cas9-NLS-6xHis (addgene plasmids#62933) and the specific sgRNA sequences were inserted successively, forming the plasmid Ptac-Cas9-T7-sgRNA as shown in Fig. S2. The generation protocols of other plasmids were described in supplementary information.

### Expression and purification of Cas9 or Cas9 RNP

The *E.coli* Rosetta (DE3) competent cells were transformed with the plasmid pET-Cas9-NLS-6xHis or Ptac-cas9-T7sgRNA to produce the corresponding Cas9 or Cas9 RNP, respectively. The selected monoclones were inoculated in Luria-Bertani (LB) broth containing 100 mg/ml of ampicillin at 37°C overnight. The cells were then diluted 1:100 into the same growth medium and grown at 37°C to OD600 0.6. Next, the culture was incubated at 4°C for 30 minutes, followed by adding isopropyl β-D-thiogalactoside (IPTG) at a final concentration of 1 mM for induction of Cas9 or Cas9 RNP at 18 °C. After 16 h, the cells were collected, re-suspended in lysis buffer (20 mM Tris-HCl. pH 8.0, 500 mM NaCl,10 mM imidazole, 10% glycerol), broken by sonication, and purified on Ni-NTA resin. The resulting Cas9 or Cas9 RNP was desalted and concentrated to ~2 mg/ml by Amicon Ultra centrifugal filters (Millipore) and stored at −20°C in storage buffer (10 mM Tris-Cl, pH 7.4, 500 mM NaCl, 0.1 mM EDTA, 1 mM DTT, 50% glycerol).

### In vitro transcription of sgRNAs

Linear DNA fragment containing the T7 promoter binding site followed by 20 bp sgRNA target sequence was transcribed in vitro using the T7 High Yield RNA Synthesis Kit (NEB) according to the manufacturer’s instructions. The resulting sgRNAs were purified using PureLink^®^ RNA Mini Kit (Thermo Fisher Scientific) and stored at -80 °C.

### Digestion of dsDNA by Cas9 RNP

To prepare Cas9 RNP in vitro by conventional method, we added Cas9 protein to the specific sgRNAs at 1:1.2 molar ratio with slowly swirling followed by incubation at 37°C for 10 min. The Cas9 RNP prepared from *E. coil* can be directly applied to digest dsDNA. The digestion was typically carried out in a 10 μl reaction mixture, composed of 1 μl 10 × buffer 3.1 (NEB), 200 ng Cas9 RNP, 300 ng plasmid, at 37°C for 30 minutes followed by termination of the reaction at 80°C for 10 minutes.

### RNA-sequencing

We extracted the incorporated sgRNAs from RNA electrophoresis gel by RNA extraction kit (NEB) and followed by two cycles of RT-PCR using primers in Balancer NGS Library Preparation Kit for small/microRNA. The library was gel-extracted on an 8% native PAGE gel and sequenced on an Illumina MiSeq.

## Acknowledgments

We thank T.A. Liu for kindly gifting the genes involving astaxanthin biosynthetic pathway. This research was supported by Key Technical Innovation Foundation of Hubei Province (NO.2017ACA174 to L.X. Ma) and National Foundation of Hubei Province (NO. 201700963 to Y. Liu).

## Author contributions

W.Q. Li and S.T. Li contributed equally to the work. W.Q. Li, Y. Liu and L.X. Ma designed the experiments; W.Q. Li and S.T. Li executed the experiments including construction and purification of Cas9 RNP, as well as in vitro assembly of DNA fragments; J. Qiao and Y. Liu executed cell culture and in vivo genome editing experiments; F. Wang and R. Y. He invented T5-assisted cloning method; W.Q. Li and Y. Liu analyzed the data; Y. Liu wrote the manuscript and all authors revised and agreed to the manuscript.

## Additional information

**Supplementary Information** accompanies this paper at XX.

## References

1. Jiang, F. & Doudna, J.A. CRISPR-Cas9 Structures and Mechanisms. Annu. Rev. Biophys. 46, 505–529 (2017).

2. Wang, H., La Russa, M. & Qi, L.S. CRISPR/Cas9 in Genome Editing and Beyond. Annu. Rev. Biochem. 85, 227–264 (2016).

3. Komor, A.C., Badran, A.H. & Liu, D.R. CRISPR-Based Technologies for the Manipulation of Eukaryotic Genomes. Cell 168, 20–36 (2017).

4. Koonin, E.V., Makarova, K.S. & Zhang, F. Diversity, classification and evolution of CRISPR-Cas systems. Curr. Opin. Microbiol. 37, 67–78 (2017).

5. Jinek, M., et al. A programmable dual-RNA-guided DNA endonuclease in adaptive bacterial immunity. Science 337, 816–821 (2012).

6. Scott, D.A. & Zhang, F. Implications of human genetic variation in CRISPR-based therapeutic genome editing. Nat. Med. 23, 1095–1101 (2017).

7. Fu, Y., Reyon, D. & Joung, J.K. Targeted genome editing in human cells using CRISPR/Cas nucleases and truncated guide RNAs. Methods. Enzymol. 546, 21–45 (2014).

8. Cho, S.W., et al. Analysis of off-target effects of CRISPR/Cas-derived RNA-guided endonucleases and nickases. Geno. Res. 24, 132–141 (2014).

9. Liang, P., Zhang, X., Chen, Y. & Huang, J. Developmental history and application of CRISPR in human disease. J. Gene. Med. 19(2017).

10. Porteus, M.H. & Carroll, D. Gene targeting using zinc finger nucleases. Nat. Biotechnology. 23, 967–973 (2005).

11. Wood, A.J., et al. Targeted genome editing across species using ZFNs and TALENs. Science 333, 307 (2011).

12. Barrangou, R. & Doudna, J.A. Applications of CRISPR technologies in research and beyond. Nat. Biotechnology. 34, 933–941 (2016).

13. Wang, T., Wei, J.J., Sabatini, D.M. & Lander, E.S. Genetic screens in human cells using the CRISPR-Cas9 system. Science 343, 80–84 (2014).

14. Shalem, O., et al. Genome-scale CRISPR-Cas9 knockout screening in human cells. Science 343, 84–87 (2014).

15. Jiang, W., et al. Cas9-Assisted Targeting of CHromosome segments CATCH enables one-step targeted cloning of large gene clusters. Nat. Commun. 6, 8101–8119 (2015).

16. Gu, W., et al. Depletion of Abundant Sequences by Hybridization (DASH): using Cas9 to remove unwanted high-abundance species in sequencing libraries and molecular counting applications. Genome. Biol. 17, 41 (2016).

17. Gao, X., et al. Treatment of autosomal dominant hearing loss by in vivo delivery of genome editing agents. Nature 553, 217–221 (2018).

18. Staahl, B.T., et al. Efficient genome editing in the mouse brain by local delivery of engineered Cas9 ribonucleoprotein complexes. Nat. Biotechnology. 35, 431–434 (2017).

19. Wang, M., et al. Efficient delivery of genome-editing proteins using bioreducible lipid nanoparticles. Proc. Natl. Acad. Sci. USA 113, 2868–2873 (2016).

20. Zuris, J.A., et al. Cationic lipid-mediated delivery of proteins enables efficient protein-based genome editing in vitro and in vivo. Nat. Biotechnology. 33, 73–80 (2015).

21. Kim, S., Kim, D., Cho, S.W., Kim, J. & Kim, J.S. Highly efficient RNA-guided genome editing in human cells via delivery of purified Cas9 ribonucleoproteins. Genome. Research. 24, 1012–1019 (2014).

22. Liu, Y., et al. In Vitro CRISPR/Cas9 System for Efficient Targeted DNA Editing. MBio 6, e01714–01715 (2015).

23. Anders, C. & Jinek, M. In vitro enzymology of Cas9. Methods. Enzymol. 546, 1–20 (2014).

24. Pingoud, A., Wilson, G.G. & Wende, W. Type II restriction endonucleases – a historical perspective and more. Nucleic. Acids. Res. 44, 7489–7527 (2016).

25. Shetty, R.P., Endy, D. & Knight, T.F., Jr. Engineering BioBrick vectors from BioBrick parts. J. Biol. Eng. 2, 5 (2008).

26. Engler, C., Gruetzner, R., Kandzia, R. & Marillonnet, S. Golden gate shuffling: a one-pot DNA shuffling method based on type IIs restriction enzymes. PLoS. One. 4, e5553–5562 (2009).

27. Cameron, D.E., Bashor, C.J. & Collins, J.J. A brief history of synthetic biology. Nat. Rev. Microbio. 12, 381–390 (2014).

28. Keasling, J.D. Synthetic biology for synthetic chemistry. ACS. Chem. Biol. 3, 64–76 (2008).

29. Li, M.Z. & Elledge, S.J. Harnessing homologous recombination in vitro to generate recombinant DNA via SLIC. Nat. Methods. 4, 251–256 (2007).

30. Stevenson, J., Krycer, J.R., Phan, L. & Brown, A.J. A practical comparison of ligation-independent cloning techniques. PLoS. One. 8, e83888 (2013).

31. Liang, J., Liu, Z., Low, X.Z., Ang, E.L. & Zhao, H. Twin-primer non-enzymatic DNA assembly: an efficient and accurate multi-part DNA assembly method. Nucleic. Acids. Res. 45, e94 (2017).

32. Hartley, J.L., Temple, G.F. & Brasch, M.A. DNA cloning using in vitro site-specific recombination. Genome. Research. 10, 1788–1795 (2000).

33. Gibson, D.G., et al. Enzymatic assembly of DNA molecules up to several hundred kilobases. Nat. Methods. 6, 343–345 (2009).

34. Gasiunas, G., Barrangou, R., Horvath, P. & Siksnys, V. Cas9-crRNA ribonucleoprotein complex mediates specific DNA cleavage for adaptive immunity in bacteria. Proc. Natl. Acad. Sci. USA. 109, E2579–2586 (2012).

35. Huang, L., et al. Efficient and specific gene knockdown by small interfering RNAs produced in bacteria. Nat. Biotechnology. 31, 350–356 (2013).

36. Qing, G., et al. Cold-shock induced high-yield protein production in *Escherichia coli*. Nat. Biotechnology. 22, 877–882 (2004).

37. Kweon, J., et al. Fusion guide RNAs for orthogonal gene manipulation with Cas9 and Cpf1. Nat. Commun. 8, 1723 (2017).

38. Danna, K. & Nathans, D. Specific cleavage of simian virus 40 DNA by restriction endonuclease of Hemophilus influenzae. Proc. Natl. Acad. Sci. USA. 68, 2913–2917 (1971).

39. Roberts, R.J., Vincze, T., Posfai, J. & Macelis, D. REBASE--restriction enzymes and DNA methyltransferases. Nucleic. Acids. Res. 33, D230–232 (2005).

40. Wang, J.W., et al. CRISPR/Cas9 nuclease cleavage combined with Gibson assembly for seamless cloning. Biotechniques 58, 161–170 (2015).

41. Cong, L., et al. Multiplex genome engineering using CRISPR/Cas systems. Science 339, 819–823 (2013).

42. Hsu, P.D., et al. DNA targeting specificity of RNA-guided Cas9 nucleases. Nat. Biotechnology. 31, 827–832 (2013).

43. Benoit, R.M., et al. Seamless Insert-Plasmid Assembly at High Efficiency and Low Cost. PLoS. One. 11, e0153158 (2016).

44. Zhu, D., et al. High-throughput cloning of human liver complete open reading frames using homologous recombination in *Escherichia coli*. Anal. Biochem. 397, 162–167 (2010).

45. Casini, A., Storch, M., Baldwin, G.S. & Ellis, T. Bricks and blueprints: methods and standards for DNA assembly. Nat. Rev. Mol. Cell Biol. 16, 568–576 (2015).

46. Anderson, J.C., et al. BglBricks: A flexible standard for biological part assembly. J. Biol. Eng. 4, 1 (2010).

47. Liu, J.K., Chen, W.H., Ren, S.X., Zhao, G.P. & Wang, J. iBrick: a new standard for iterative assembly of biological parts with homing endonucleases. PLoS. One. 9, e110852 (2014).

48. Ma, T., et al. Genome mining of astaxanthin biosynthetic genes from Sphingomonas sp. ATCC 55669 for heterologous overproduction in *Escherichia coli*. Biotechnol. J. 11, 228–237 (2016).

49. Li, S.Y., Zhao, G.P. & Wang, J. C-Brick: A New Standard for Assembly of Biological Parts Using Cpf1. ACS. Synth. Biol. 5, 1383–1388 (2016).

50. DeWitt, M.A., Corn, J.E. & Carroll, D. Genome editing via delivery of Cas9 ribonucleoprotein. Methods. 121–122, 9–15 (2017).

51. Harrington, L.B., et al. A thermostable Cas9 with increased lifetime in human plasma. Nat. Commun. 8, 1424 (2017).

52. Zetsche, B., et al. Cpf1 is a single RNA-guided endonuclease of a class 2 CRISPR-Cas system. Cell 163, 759–771 (2015).

53. Abudayyeh, O.O., et al. RNA targeting with CRISPR-Cas13. Nature 550, 280–284 (2017).

54. Cox, D.B.T., et al. RNA editing with CRISPR-Cas13. Science 358, 1019–1027 (2017).

